# BARe-seq enables high-throughput dissection of cis-regulatory control of transcriptional bursting

**DOI:** 10.64898/2026.08.19.745405

**Authors:** Franziska K. Lorbeer, Reyna Edith Rosales Alvarez, Katharina Bergauer, Dominic Grün, Alexander Stark

## Abstract

Transcriptional bursts determine RNA output through two kinetic parameters: burst size and burst frequency. How cis-regulatory DNA encodes these kinetic parameters remains unclear, in part because existing approaches do not combine scalable sequence perturbation with allele-resolved burst inference. Here, we developed Bulk Allele Resolution Sequencing (BARe-seq), an allele-resolved massively parallel reporter assay that enables inference of transcriptional burst parameters from bulk sequencing. Applying BARe-seq to libraries of 1000 promoters and 1000 enhancers in *Drosophila* S2 cells revealed distinct kinetic properties of promoters and enhancers. Promoter-dependent mean expression was driven by both burst size and burst frequency: TATA-box promoters showed larger bursts, whereas DPE promoters showed higher burst frequency. In contrast, enhancer strength was primarily driven by burst frequency, although specific transcription factor motifs were also associated with burst size. Thus, BARe-seq dissects cis-regulatory control of transcriptional bursting and extends allele-resolved measurements to pooled reporter assays in bulk sequencing experiments.

## Introduction

Transcription occurs in discontinuous bursts of activity, separated by inactive intervals (Elowitz et al., 2002; Golding et al., 2005). While individual bursts are driven by stochastic molecular interactions, genes exhibit reproducible steady-state bursting kinetics, indicating that these kinetics are genetically regulated rather than purely random (Garcia et al., 2013; Sanchez & Golding, 2013; Bothma et al., 2014; Lim et al., 2018; Larsson et al., 2019).

The transcriptional output is determined by both the number of transcripts produced per active period (burst size) and the occurrence rate of these periods (burst frequency) (reviewed in Nicolas et al., 2017; Raj & van Oudenaarden, 2008). While regulatory inputs modulate gene expression by changing one or both of these parameters, it remains unclear how specific cis-regulatory DNA sequences are linked to burst kinetics. Previous work has suggested a division between promoter and enhancer contributions. Promoter architecture has been primarily associated with burst size, including links between TATA-containing promoters and larger, but not more frequent, bursts (Hornung et al., 2012; Larsson et al., 2019; Pimmett et al., 2021), while enhancer activity is associated primarily with burst frequency (Falo-Sanjuan et al., 2019; Fukaya et al., 2016; Larsson et al., 2019; Hoppe et al., 2020).

Yet, perturbations of individual promoter motifs (Yokoshi et al., 2022), as well as changes in signaling, transcription factor abundance, residence time or binding affinity can affect both burst size and frequency (Senecal et al., 2014; Ochiai et al., 2020). These observations indicate that the cis-regulatory encoding of burst kinetics extends beyond a simple model in which promoters determine burst size and enhancers determine burst frequency. However, it remains underexplored how promoter and enhancer sequences—particularly their motif compositions—relate to bursting. Resolving this complexity requires assays that can evaluate whether regulatory sequence differences are reflected in burst size, burst frequency, or both.

Defining general rules that link cis-regulatory sequence to burst kinetics requires scaling these measurements. Existing methods address different aspects of this problem but fail to combine burst-parameter inference with systematic sequence dissection: Imaging-based approaches measure transcription dynamics directly but are limited in throughput and typically require dedicated reporter engineering (reviewed in Nicolas et al., 2017). Allele-resolved single-cell RNA sequencing (scRNA-seq) has enabled transcriptome-wide inference of bursting in endogenous contexts (Larsson et al., 2019, 2021; Ochiai et al., 2020), but systematic cis-regulatory perturbations remain difficult to test at endogenous loci. Conversely, massively parallel reporter assays (MPRAs) offer the scalability to assess thousands of sequences, including perturbed and designed synthetic sequences (Arnold et al., 2013; Kwasnieski et al., 2014; Arnold et al., 2017; van Arensbergen et al., 2017; Bergman et al., 2022; de Almeida et al., 2022). However, standard MPRAs quantify aggregate reporter expression and lack the allelic resolution required to infer kinetic burst parameters.

To address this methodological gap, we developed Bulk Allele-Resolution sequencing (BARe-seq). BARe-seq uses plasmid barcodes to achieve single-allele resolution from a bulk experiment. By tracking of transcripts back to uniquely barcoded plasmids, the assay isolates independent transcriptional activity of individual template copies within a pooled sample. We applied this strategy for high-throughput inference of burst parameters across large libraries of cis-regulatory elements and quantified burst parameters across libraries of 1000 promoters and 1000 enhancers in *Drosophila* S2 cells. Using BARe-seq, we recovered established relationships between promoter motifs and bursting, including the association of TATA-containing promoters with larger bursts, and identified additional motif relationships involving DPE (downstream promoter element), Inr (Initiator), E-box and Ohler6 motifs. We further showed that promoters and enhancers contribute asymmetrically to transcriptional bursting: promoter-dependent mean expression variation is globally driven by both burst size and frequency changes, while enhancer-dependent variation is primarily driven by burst frequency changes. Yet, beneath this global trend, we found that specific transcription factor motifs in enhancers are associated with burst size and frequency and used designed enhancer variants to test how motif identities and the enhancer-sequence context contribute to bursting. Together, our results establish BARe-seq as a scalable approach for dissecting how cis-regulatory sequences encode transcriptional bursting kinetics.

## Results

### BARe-seq uses plasmid barcoding and transcript UMIs to count the mRNA output per plasmid

BARe-seq uses high-complexity plasmid barcodes to recover transcript counts from individual reporter plasmids in bulk sequencing experiments. By introducing random DNA barcodes (plasmid-specific unique molecular identifiers = pUMIs) into pooled libraries of regulatory elements, we generated a library of effectively uniquely barcoded plasmids, allowing each expressed RNA molecule to be assigned to its plasmid of origin. During sample processing, each RNA molecule is additionally associated with a transcript unique molecular identifier (tUMI) to ensure exact counting of the mRNA molecules produced. Consequently, each regulatory element yields a distribution of transcript counts across its associated plasmids, and this count distribution enables the inference of transcriptional burst parameters (Figure 1A). BARe-seq thus achieves allele-resolved mRNA quantification from bulk sequencing experiments. By relying on a pooled measurement, the assay circumvents technical sources of cell-to-cell variability inherent to scRNA-seq experiments that otherwise complicate the inference of biological noise (Grün et al., 2014; Rosales-Alvarez et al., 2023).

**Figure 1:**
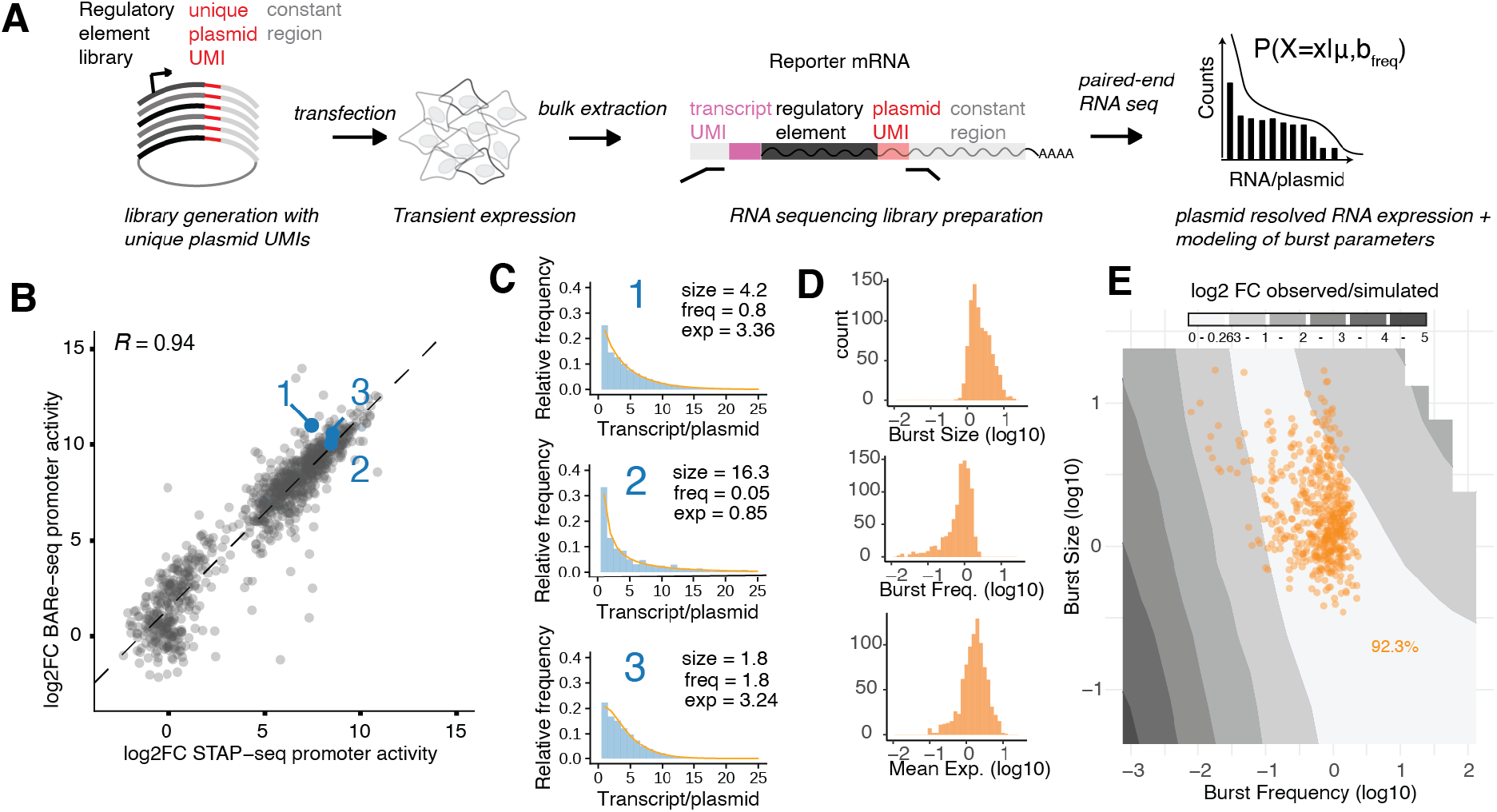
BARe-seq enables high-throughput burst inference from allele-resolved bulk measurements. (A) Schematic showing the workflow of a BARe-seq experiment. After generation and labeling of a regulatory element library with plasmid unique molecular identifiers (pUMIs), plasmids are transfected and transiently expressed in *Drosophila* S2 cells. Following bulk RNA extraction, the RNA is processed and labeled with transcript UMIs (tUMIs). Following paired-end next-generation sequencing, pUMI and tUMI information is integrated to obtain per-plasmid transcript count distributions for each regulatory element. (B) Scatter plot comparing the BARe-seq derived promoter activity with previously published STAP-seq promoter activity (Haberle et al., 2019) *R* = Pearson’s *R*. Three data points are highlighted, for which per-plasmid transcript frequency distributions are shown as blue histograms in (C). Negative binomial fits of the distributions are overlaid in orange. (D) Histograms of all inferred parameters, displaying the dynamic ranges of burst size, burst frequency, and mean expression. (E) Contour plot of the simulated parameter space, showing burst size as a function of burst frequency on a log_10_ scale. Greyscale indicates the distance of inferred parameters from the ground truth as absolute log_2_ fold-change (|log2FC|), where the white region (range 0 – 0.263) exhibits the highest inference accuracy. Data points of inferred parameters of promoter library are projected on the contour plot with 92.3% of data points (orange dots) falling into the area of highest confidence.

### The promoter BARe-seq library is labeled with effectively unique pUMIs

To establish BARe-seq in the context of promoters, we selected 1000 deeply characterized *Drosophila* S2 cell promoters spanning a broad range of activities measured by the promoter activity reporter assay self-transcribing active core promoter sequencing (STAP-seq) (see Methods) (Haberle et al., 2019). We cloned these promoters as a promoter-BARe-seq library in which each plasmid carried a 20-nucleotide random barcode that served as a pUMI (Figure 1A, Supplemental Figure 1A). Next-generation sequencing confirmed the high complexity of the plasmid pool: among 10^7^ sequenced promoter – pUMI combinations, 98.3% were unique, making barcode collision within the assay highly unlikely so that the pUMIs successfully resolve single plasmids (Supplemental Figure 1B).

### BARe-seq recovers promoter activities and produces reproducible per-plasmid expression distributions

We next asked whether BARe-seq accurately recapitulates prior measurements of promoter activity (Arnold et al., 2017; Haberle et al., 2019). We placed the promoter-BARe-seq library under the control of the highly active *zfh1* enhancer, transfected it into *Drosophila* S2 cells, and quantified reporter expression by bulk next-generation sequencing (see Methods). At the aggregate level – by collapsing reads across all pUMIs – BARe-seq activity scores were highly correlated with previously published STAP-seq data (Pearson’s correlation coefficient *R* = 0.94) (Haberle et al., 2019). This confirmed that pUMI insertion did not disrupt reporter performance and that BARe-seq captured promoter activities (Figure 1B). Concurrently, because each sequencing read contained the promoter identity, pUMI, and tUMI, reporter RNA molecules could be accurately counted at the level of individual plasmids (Figure 1A, Supplemental Figure 1C). This yielded a distribution of reporter transcripts per plasmid for each promoter.

These distributions differed between different promoters and were reproducible across replicates (Figure 1C, Supplemental Figure 1D). Thus, BARe-seq simultaneously recovers conventional aggregate reporter readouts and provides the plasmid-resolved transcript distributions needed for burst parameter inference.

### Burst parameters can be inferred from per-plasmid transcript distributions using the two-state model of stochastic transcription

The dynamics of gene expression are captured by the two-state model of stochastic transcription described by Peccoud and Ycart (1995) (Peccoud & Ycart, 1995) (Supplemental Figure 2A). Assuming predominant bursting behavior, the stationary state of this model is described by a negative binomial distribution with parameters mean *μ* and dispersion *r* (Raj et al., 2006; Grün et al., 2014). In this limit, dispersion *r* equates to burst frequency, while the ratio of *μ/r* defines burst size. To ensure accurate model fitting, we applied a Kolmogorov-Smirnov (KS) goodness-of-fit threshold (*p* > 0.01) between empirical and inferred distributions, retaining data for 547 promoters from replicate 1 and 593 promoters from replicate 2. The inferred parameters proved reproducible across independent experimental replicates (*R* = 0.82 for burst size, *R* = 0.66 for burst frequency, and *R* = 0.75 for mean expression) (Supplemental Figure 2B). To further improve the robustness and reproducibility, we pooled count data from two replicates and inferred the burst parameters for 608 promoters. We further restricted downstream analyses to promoters above an expression threshold guided by the observed bimodal distribution of means, where parameter estimates are more robust (*R* = 0.92 for burst size, *R* = 0.74 for burst frequency, and *R* = 0.9 for mean expression) resulting in a total of 492 promoters with confidently inferred burst parameters (Supplemental Figure 2C-E). The inferred parameters captured a broad dynamic range, with burst size, frequency and mean expression each spanning nearly two orders of magnitude (Figure 1D).

### Simulations identify the parameter regime for accurate inference

To verify the parameter space in which burst parameters can be accurately inferred with our modeling approach, we designed a simulation experiment. We generated count distributions across a wide range of burst frequencies and mean expressions (and, hence, burst sizes), and applied our inference pipeline to compare the inferred values to the known ground truth. This allowed us to define a high-accuracy region where the inferred values deviate from the ground truth by no more than 20% (log2FC < 0.263) (Figure 1E, white region). From our experimental promoter data, 92.3% of the inferred parameters fell within this high-accuracy region (Figure 1E).

### BARe-seq recovers the established association between TATA box-containing promoters and larger bursts

We first asked whether BARe-seq recovers established relationships between promoter sequence and burst kinetics. Prior studies in yeast, *Drosophila* and mammalian cells have consistently linked TATA box-containing promoters to increased burst size (Hornung et al., 2012; Larsson et al., 2019; Pimmett et al., 2021; Yokoshi et al., 2022). Our promoter library represents characteristic *Drosophila* promoter types in terms of core promoter motifs and motif combinations, including TATA box-containing promoters (Figure 2A, Supplemental Figure 3A). Consistent with previous work, TATA box-containing promoters associated with higher burst sizes and lower burst frequencies than promoters without a TATA box. This relationship persisted even when accounting for differences in mean expression (Supplemental Figure 3B).

**Figure 2:**
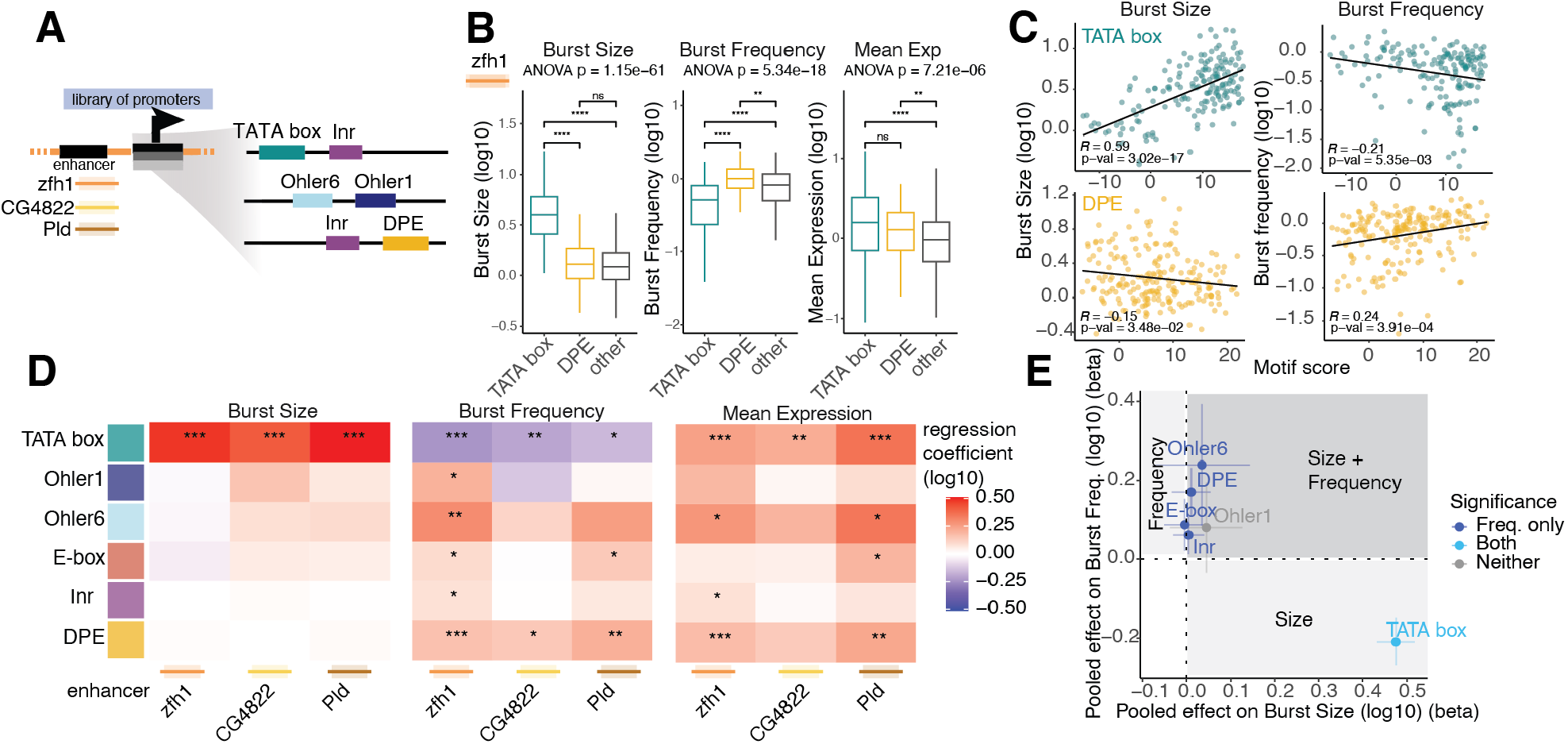
Promoter motifs are associated with differences in both burst size and frequency. (A) *Drosophila* core promoters occur in characteristic combinations, which are represented in the promoter-BARe-seq library. (B) Three-way analysis of TATA box, DPE and other promoters (“other”) for burst size, burst frequency and mean expression. Significance was determined via ANOVA followed by Tukey’s post-hoc test. (C) Scatter plots showing burst size (left) or burst frequency (right) as a function of TATA box (top) and DPE (bottom) motif scores (D) Heatmap of motif-associated regression coefficients from multivariable models of promoter-BARe-seq data in the zfh1, CG4822, and Pld contexts. Coefficients are shown on the log_10_-transformed scale. Significance markings: * *p* < 0.05, ** *p* < 0.01, *** *p* < 0.001. (E) Pooled fixed-effect meta-analysis estimates for the association between motif presence and burst size (bsize, x-axis) or burst frequency (bfreq, y-axis); points represent pooled log_10_-scale regression coefficients, error bars indicate 95% confidence intervals, and colored points denote motifs significant for either outcome at *p*<0.05.

### TATA and DPE show distinct motif-presence and motif-strength relationships with burst parameters

To extend the analysis, we compared the TATA box against the downstream promoter element (DPE). Both are prominent *Drosophila* developmental core promoter motifs but occur in a largely mutually exclusive manner (Ohler, 2006; Arnold et al., 2017). As expected, the TATA box-containing promoters associated with higher burst size and lower frequency, while DPE-containing promoters showed higher burst frequencies but no significant difference in burst size (Figure 2B). Having observed these motif associations, we next asked whether motif affinity, approximated by the position weight matrix (PWM) score, was quantitatively related to burst parameters. The TATA box PWM score correlated positively with burst size (*R* = 0.59, *p* = 3.02*10^-17^) and negatively with burst frequency (*R* = -0.21, *p* = 0.0054). Conversely, the DPE PWM score correlated positively with burst frequency (*R* = 0.24, *p* = 0.0004) and negatively with burst size (*R* = -0.15, *p* = 0.035) (Figure 2C). Therefore, BARe-seq resolves motif-burst relationships down to the quantitative level of motif affinity.

### Promoter motif–burst relationships generalize across enhancer contexts

To determine whether the relationships between promoter types and burst parameters persist across enhancer contexts, we performed the promoter-BARe-seq screen with two additional enhancers (CG4822 and Pld). These enhancers were selected based on their activity in a previous self-transcribing active regulatory region sequencing (STARR-seq) screen (Zabidi et al., 2015) and confirmed by luciferase assays (Supplemental Figure 3C,D). We inferred burst parameters for each dataset separately and subsequently applied multivariable regression to assess motif associations with the parameters while accounting for motif co-occurrence (Figure 2D, Supplemental Figure 3E). Across the three enhancer contexts, the TATA box consistently associated with higher burst size and lower burst frequency, while the DPE motif associated with higher burst frequency. In contrast, E-box, Inr and Ohler6 associated significantly with higher burst frequency only in individual enhancer contexts, suggesting their kinetic contributions may depend on enhancer mediated mechanisms. To summarize global motif effects, we integrated the regression coefficients and standard errors by meta-analysis (Figure 2E). This analysis identified the TATA box as the only significant positive predictor of burst size, while DPE, E-box, Inr and Ohler6 motifs were significant positive predictors of burst frequency. Therefore, the core promoter motifs encode distinct but generalizable kinetic features that are robust across enhancer contexts.

### Promoter-dependent expression variation is driven by both burst size and frequency

Having validated promoter-BARe-seq by recovering kinetic signatures for core promoter motifs, we next asked how the interplay between promoters and enhancers regulates bursting. The functional interactions between the two regulatory elements are central to gene regulation, but it remains unclear how their respective contributions to transcription are driven by burst size and burst frequency. BARe-seq allowed us to decouple the effects of promoters and enhancers by varying one element while holding the other constant. We first analyzed promoter-dependent variation in expression in the three promoter-BARe-seq screens described above (Figure 2), which assess the 1000-promoter library against three constant enhancers. We ordered the promoters by mean expression from weak to strong and observed that both burst size and burst frequency increased with mean expression, a trend that was true for all three enhancer contexts (Figure 3A). Linear model slopes were positive for both parameters across all three contexts (Figure 3C), indicating that promoter-dependent differences in expression are driven by changes in both burst size and burst frequency.

**Figure 3:**
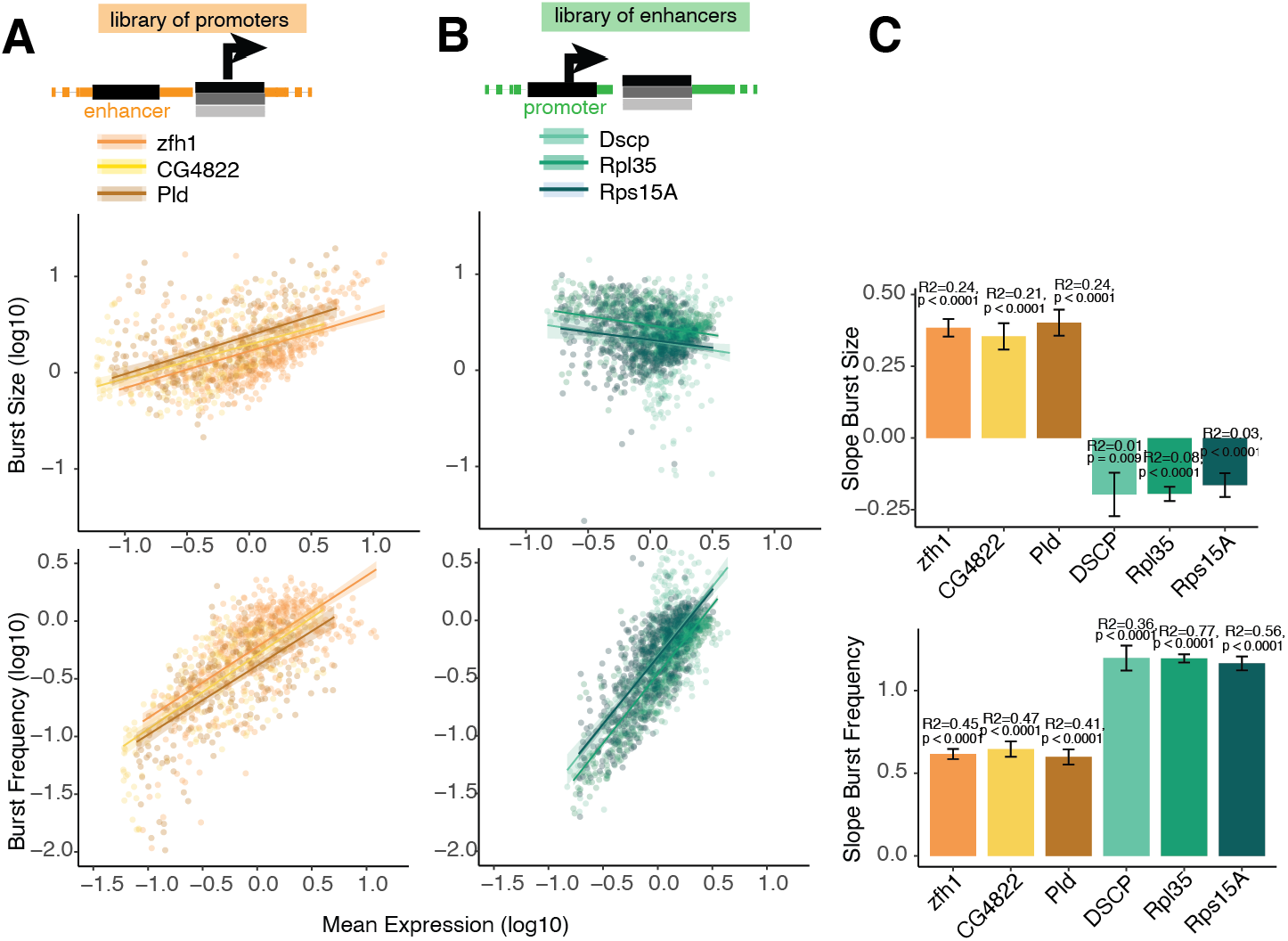
Promoter-dependent variation involves both burst size and frequency, while enhancer-dependent variation primarily reflects burst frequency. (A-B) Scatter plots showing the relationships between burst size and mean expression (left column) and burst frequency and mean expression (right column) for the promoter libraries (A) and the enhancer libraries (B). Solid lines represent linear regression fits, with shaded areas indicating the 95% confidence intervals. Axes were limited for better visibility, points outside the ranges were clipped. (C) Bar graphs of slopes with error bars representing the 95% confidence intervals and *p*-values of the log_10_ transformed parameters indicated.

### Enhancer-dependent expression variation is reflected primarily in burst frequency

To assess enhancer-dependent variation in expression, we inverted the experimental design by coupling a constant promoter to a library of 1000 well-characterized enhancers, adapting the STARR-seq design (Arnold et al., 2013; de Almeida et al., 2022). We measured enhancer activity with DSCP, a synthetic developmental core promoter, and additionally with two highly active housekeeping-type promoters, Rpl35 and Rps15a (Supplemental Figure 4A)(Pfeiffer et al., 2008; Arnold et al., 2013; Zabidi et al., 2015). Aggregate enhancer-BARe-seq enhancer activity correlated well with the previously published STARR-seq measurements (*R* = 0.84) (de Almeida et al., 2022) (Supplemental Figure 4B). After pooling replicates and applying the same quality thresholds used for the promoter analysis, burst parameters were inferred for 451 enhancers with DSCP, 685 with Rpl35 and 601 with Rps15a (*KS p-value* > 0.01 and mean expression threshold; Supplemental Figure 4C). The enhancers showed a wide range of mean expression and burst parameters comparable to the promoters (Supplemental Figure 4D), allowing us to order enhancers by mean expression from weak to strong. In contrast to promoters, only burst frequency but not burst size increased with enhancer strength (mean expression). Linear model slopes were only positive for burst frequency but weakly negative for burst size (Figure 3B,C). Thus, enhancer-dependent differences in mean expression were driven primarily by burst frequency, not burst size. Together, these data indicate that promoters and enhancers contribute asymmetrically to burst kinetics. Across promoters, mean expression was driven by both burst size and frequency, yet across enhancers, mean expression changes were primarily controlled by changes in burst frequency.

### Enhancer motif composition is associated with distinct bursting regimes

Although enhancer-dependent variation in mean expression is driven primarily by burst frequency, we nevertheless observed distinct variation in burst size across the enhancer library (Figure 3B). We hypothesized that specific sequence features within enhancers can account for the burst size variation. Because enhancers contain complex combinations of transcription factor motifs, individual motif types could associate with different burst parameters. To test this, we compared burst parameters between enhancers as a function of the enhancers’ motif content, focusing on the ten motifs previously identified as most important for enhancer activity in *Drosophila* S2 cells (Arnold et al., 2013; Yáñez-Cuna et al., 2014; de Almeida et al., 2022) (Figure 4A). To account for patterns of motif co-occurrence, we used multivariable regression analysis and assessed associations of individual motifs with burst size and burst frequency in each promoter context (Figure 4B, Supplemental Figure 5A-C). Several enhancer motifs, including ETS, AP-1 and SREBP, were significantly associated with higher burst size in at least one promoter context. Thus, although enhancer-dependent variation in mean expression is driven primarily by burst frequency, enhancer motif composition is also associated with differences in burst size.

**Figure 4:**
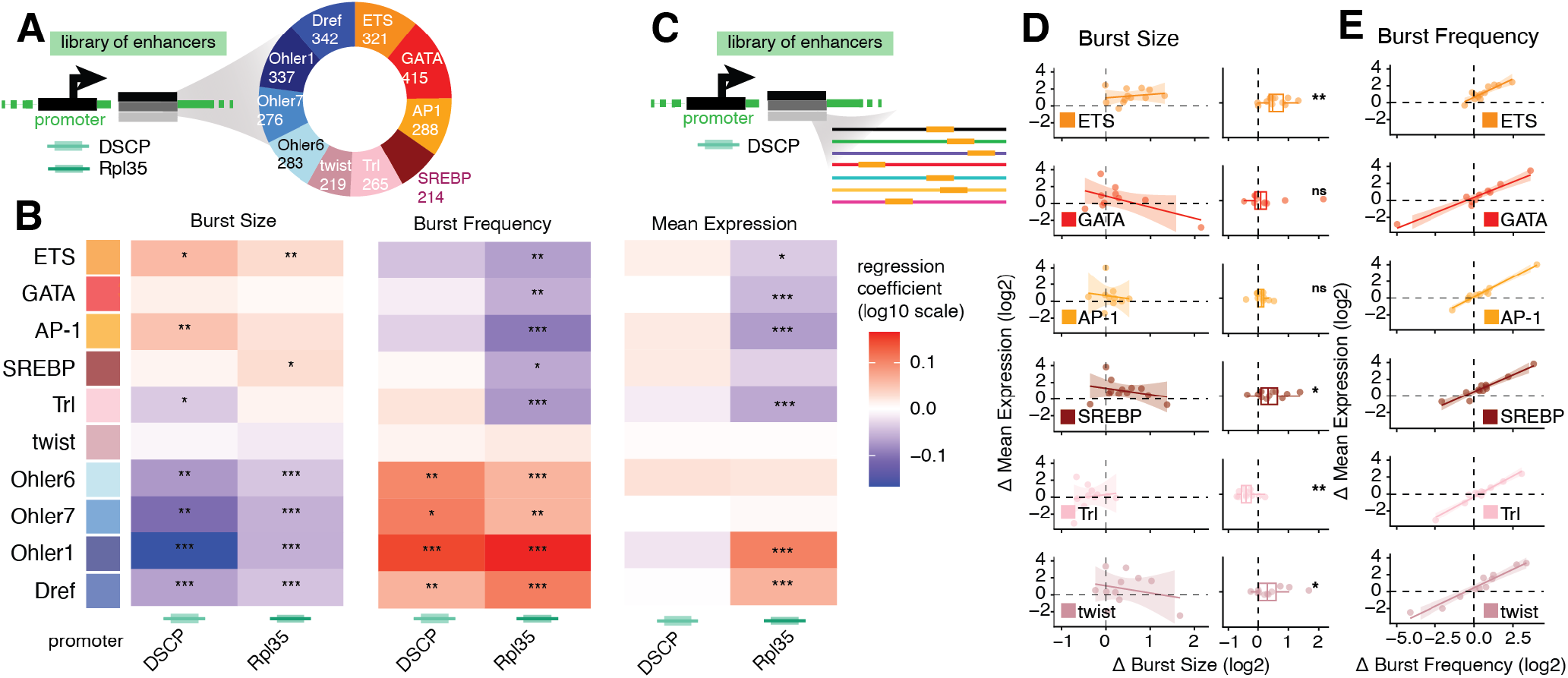
Enhancer motifs are associated with distinct burst size and frequency patterns. (A) Donut plot showing the distribution of the 10 major S2 cell enhancer motifs. (B) Heatmap showing motif-associated regression coefficients from multivariable regression models of enhancer-BARe-seq data in the DSCP and Rpl35 promoter contexts. Coefficients are shown on the log_10_-transformed outcome scale. (C) Schematic of the experimental set up for testing motif influence of bursting by pasting individual motif instances into neutral enhancer regions. (D,E) Scatter plots with linear regression lines of relative burst size (D) and relative burst frequency (E) versus the relative mean expression of pasted developmental motifs. Relative parameters (Δ) were calculated by comparing the inferred values of the inserted motif sequences against their matched wild-type base enhancers. Boxplots in (D) show relative burst sizes, with significant changes indicated (paired Wilcoxon rank-sum test, significance markings: * *p* < 0.05, ** *p* < 0.01, *** *p* < 0.001)

### Synthetic motif insertions probe the causal effects of individual motifs on bursting

While multivariable regression establishes motif associations, sequence perturbations are used to test causality. Previous studies have shown that motif deletions and insertions can alter enhancer activity in S2 cells (Yáñez-Cuna et al., 2014; de Almeida et al., 2022). To determine whether adding specific enhancer motifs alters mean expression via changes in burst size, frequency, or both, we generated synthetic enhancer variants by inserting single motif copies into existing enhancers at positions predicted to be not essential for enhancer activity (non-disruptive positions) (Methods and Figure 4C, Supplemental Figure 5D)). We obtained results from 15 wild-type enhancers spanning a range of burst sizes, burst frequencies and mean expression values (Supplemental Figure 5E). Replicate measurements of the insertion constructs were well correlated (N=2; burst size: *R* = 0.89, burst frequency: *R* = 0.71, mean expression: *R* = 0.9, Supplemental Figure 5F) and allowed burst parameter inference on average for 73% of the enhancers per motif. For each motif, we then compared mean expression, burst size and burst frequency for each enhancer before and after motif insertion, and then grouped relative changes in mean expression, burst size, and burst frequency by the type of inserted motif.

### Motif insertions alter burst size independently of enhancer-activity changes

Adding individual motif copies changed enhancer mean expression for all motifs (Fig. 4D, E). To determine the mechanistic basis for the changes in mean expression (Δ mean expression) we assessed the correlation with Δ burst size and Δ burst frequency. The Δ mean expression was well correlated with Δ burst frequency (average slope = 0.86, all *p* < 0.01), but not with Δ burst size (average slope = -0.42, not significant; Figure 4D-E; Supplemental Figure 5G). This is consistent with the global analysis above, in which enhancer-dependent variation in mean expression was determined primarily by burst frequency. However, motif insertions nevertheless produced motif-specific changes in burst size: ETS, SREBP and twist motif insertions increased burst size, while Trl motif insertion decreased it; GATA and AP-1 motifs had no significant effect (Figure 4D, boxplots). These results reconcile the global and motif-level enhancer analyses: changes in enhancer mean expression are driven by burst frequency, whereas specific enhancer motifs can additionally tune burst size.

## Discussion

BARe-seq combines the scalability and sequence perturbability of MPRAs with the allelic resolution required to infer transcriptional burst parameters. In doing so, it bridges the gap between imaging-based measurements of transcriptional dynamics, allele-resolved scRNA-seq, and conventional aggregate reporter assays. Because BARe-seq extracts allele-resolved count distributions from a pooled bulk sample, it bypasses the technical noise and cell-to-cell variability of scRNA-seq experiments (Rosales-Alvarez et al., 2023). By overcoming these technical bottlenecks, we were able to map cis-regulatory sequence to function for burst size and frequency in high-throughput.

Applying BARe-seq to the systematic analysis of promoter-enhancer combinations revealed an asymmetry between promoters and enhancers. Across promoter libraries, mean expression correlated with both burst size and burst frequency. Across enhancer libraries, mean expression varied predominantly with burst frequency. Promoter motif composition was also associated with burst parameters across different enhancer contexts: the presence of a TATA box was linked with higher burst size, while DPE, E-box, Inr and Ohler6 were associated with higher burst frequency. These sequence dependencies extend previous observations from individual *Drosophila* and human promoters that linked the DPE or Inr motif to elevated burst frequency (Hendy et al., 2017; Yokoshi et al., 2022) and are consistent with biochemical studies showing how core promoters differentially engage the core transcriptional machinery (Serebreni et al., 2023). For example, TAF1 (TATA-associated factor 1), a TFIID subunit implicated in the negative regulation of burst size, associates more strongly with DPE promoters (Pennington et al., 2013; Serebreni et al., 2023). This biochemical bias is in line with our observation that DPE promoters exhibit an inverse kinetic signature to TATA box-containing promoters linking them to higher burst frequency and lower burst size. Our enhancer analyses clarify that enhancer function cannot be reduced to burst frequency alone. At the level of overall enhancer activity, differences in mean expression tracked primarily with burst frequency, consistent with previous studies (Falo-Sanjuan et al., 2019; Fukaya et al., 2016; Hoppe et al., 2020). However, at the level of motif composition, enhancer sequence content was also associated with burst size, and designed motif insertions revealed motif-specific changes in burst size. In these perturbation experiments, changes in mean expression were reflected mainly in burst frequency, but ETS, SREBP, twist and Trl insertions produced distinct burst-size changes. These results reconcile the global enhancer-frequency relationship with motif-level burst-size effects: Burst frequency may reflect the integrated output of the enhancer sequence as a whole, including cooperative or context-dependent interactions among motifs, rather than the contribution of any single motif class. This distinction aligns with the observation that different transcription factors binding the enhancer either affect burst frequency or both parameters simultaneously (Keller et al., 2020). Furthermore, this may explain why previous reports have linked signaling inputs which affected enhancer activity to both burst size and burst frequency (Lee et al., 2019; Ochiai et al., 2020; Senecal et al., 2014).

As an MPRA, BARe-seq simplifies regulatory complexity and enables testing of sequence logic. Each construct tests one enhancer-promoter pair at a fixed distance, allowing sequence effects to be compared independent of the endogenous chromatin context. Burst parameters are inferred from sequencing-derived steady-state distributions rather than measured directly in live cells. The inference therefore depends on the assumptions of a simplified two-state model (Raj et al., 2006), even though additional promoter states and more complex kinetic schemes have been proposed (Tunnacliffe & Chubb, 2020).

Ultimately, the scalability and conceptual simplicity of BARe-seq make it transferable to other biological questions and assays. By applying this strategy to transcriptional bursting, we demonstrated that resolving transcript counts on a per-template basis allows the high-throughput mathematical separation of burst size from burst frequency. The method’s core innovation – achieving allele resolution from a bulk experiment via unique barcoding – is broadly applicable to other plasmid-based assays and questions that require single-allele resolution.

## Acknowledgements

We are grateful to Anton Larsson and Rickard Sandberg for help at the start of the project. We thank Yoav Voichek, John Boyle, and Bernardo Almeida and the remainder of the Stark lab for helpful discussions. FL received an EMBO postdoctoral fellowship (ALTF391-2021) and is supported by an ESPRIT fellowship of the Austrian Science Fund (FWF, 10.55776/ESP9305624). DG was supported by the ERC (818846 — ImmuNiche — ERC-2018-COG), by the Bundesministerium für Bildung und Forschung (BMBF) (TissueNet - 031L0311A), and by the Deutsche Forschungsgemeinschaft (DFG, German research Foundation) – SFB 1583/1 Project number: INST 93/1121-1, and SFB TRR 221 Project number: INST 89/681-1. Research in the Stark group is supported by the Austrian Science Fund (FWF, 10.55776/P36971, 10.55776/PAT3564423) and the Vienna Science and Technology Fund (WWTF, 10.47379/LS24012). Basic research at the IMP is supported by Boehringer Ingelheim GmbH and the Austrian Research Promotion Agency (FFG, FO999902549). For the purpose of open access, the author has applied a CC BY public copyright license to any Author Accepted Manuscript version arising from this submission.

## Competing Interests

DG serves on the scientific advisory board of Gordian Biotechnology.

## Author Contributions

FKL and AS conceived and developed BARe-seq and RRA and DG the mathematical modeling framework. FKL performed all experiments with the help of KB. RRA performed the mathematical modeling. FKL and RRA plotted and analyzed data, FKL generated final figures. FKL, RRA, DG and AS wrote the manuscript. FKL, DG and AS acquired funding. DG and AS supervised the study.

---

**Supplemental Figure 1:**
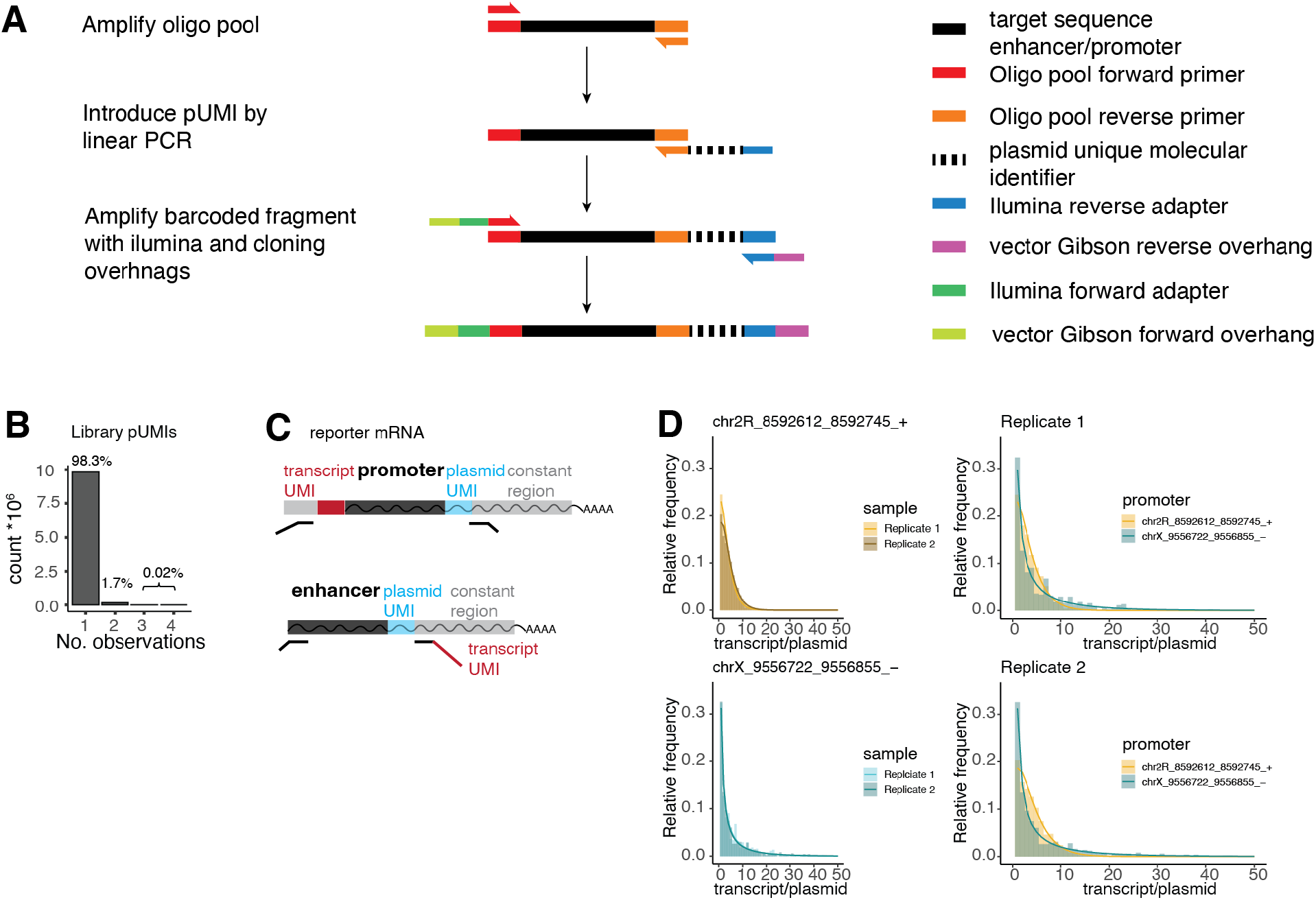
Additional information about BARe-seq library generation. (A) Workflow schematic depicting the cloning steps to generate a BARe-seq library. (B) Quality control assessing the complexity of the promoter-BARe-seq plasmid library: Barplot showing the number of observations of unique regulatory element and pUMI combinations in 10^7^ sequencing reads of the library. (C) Schematic outlining the information contained in next-generation sequencing reads for promoter and enhancer BARe-seq assay. (D) Histograms of raw BARe-seq data demonstrating reproducibility between identical regulatory elements across two replicates (left) and displaying differences between two distinct sequences within the same replicate (right). The x-axis was truncated at 50 transcripts/plasmid for visual clarity.

**Supplemental Figure 2:**
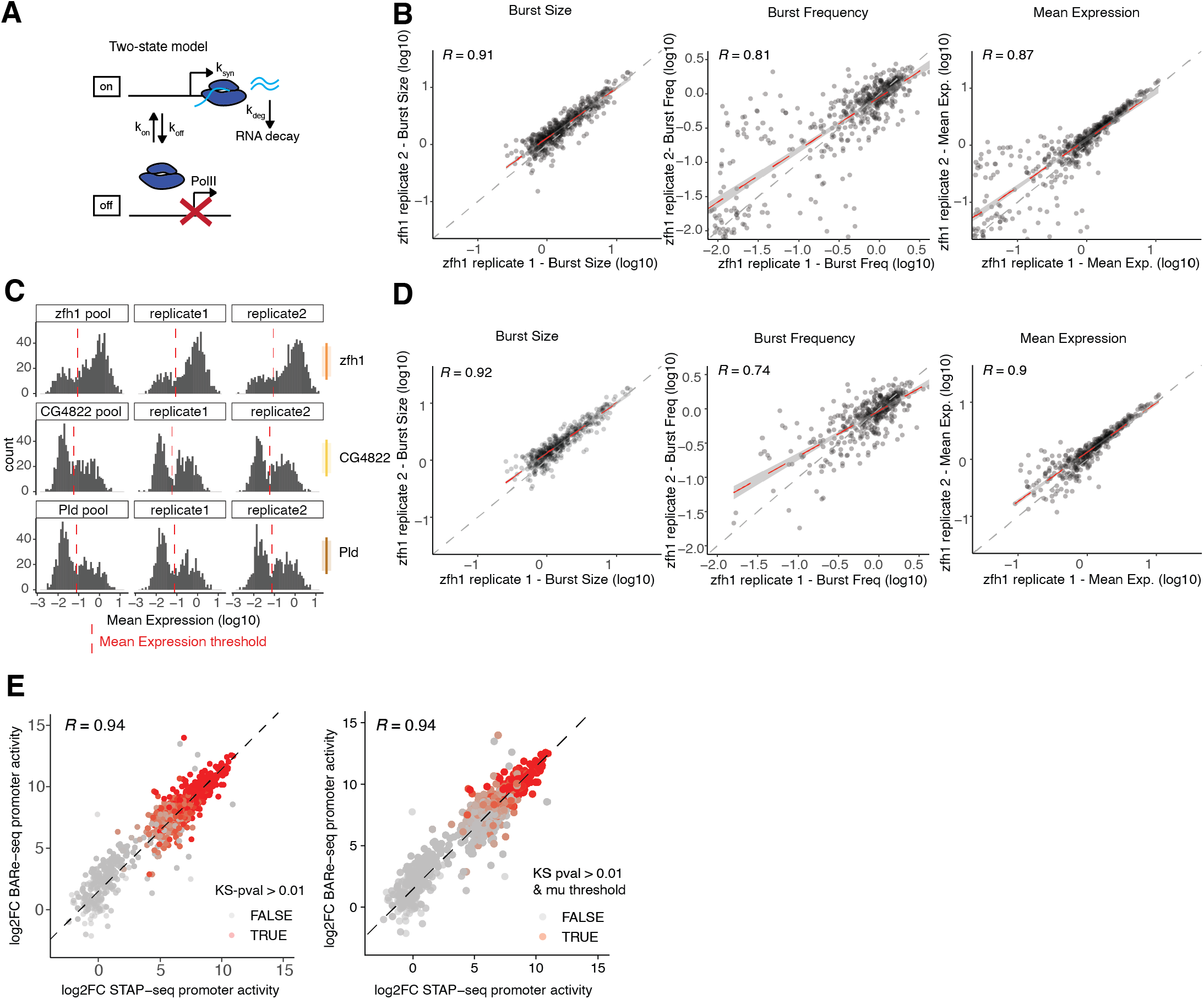
Modeling burst parameters using the two-state model of transcription. (A) Two-state model of stochastic transcription where a gene switches between active and inactive states with kinetic constants k_on_ and k_off_, respectively. The gene is transcribed only in the active state with a constant mRNA synthesis rate k_syn_, and the transcript degradation is determined by the degradation constant k_deg_. (B) Scatter plots displaying the correlation of log_10_-transformed burst size, burst frequency, and mean expression between replicates after Kolmogorov-Smirnov filtering threshold (*KS-test p* > 0.01). Red dashed line indicates the linear regression model with 95% confidence interval; dark grey dashed line indicates perfect correlation (y = x). Pearson’s R values are shown above each plot. Axes were limited for better visibility, points outside the ranges were clipped. (C) Histograms showing the bimodal distribution of inferred mean expression across different samples with no prior filtering. Red dashed lines indicate the density trough used as a threshold to remove sequences with very low transcriptional activity (see Methods). (D) Scatter plots displaying replicate correlations of log_10_-transformed burst size, burst frequency, and mean expression after applying both the KS test (*p* > 0.01) and mean expression thresholds. Red dashed line = linear regression model with standard error, dark grey dashed line = perfect correlation (y=x). Pearson’s R values are shown above each plot. (E) Scatterplot comparing the log_2_ FC activity of promoter-BARe-seq against previously published promoter activity (Haberle et al. 2019), highlighting all promoters with successfully inferred parameters after pooling meeting cut-off criteria in red (left: *KS test p* > 0.01, right: *KS test p* > 0.01 & μ threshold).

**Supplemental Figure 3:**
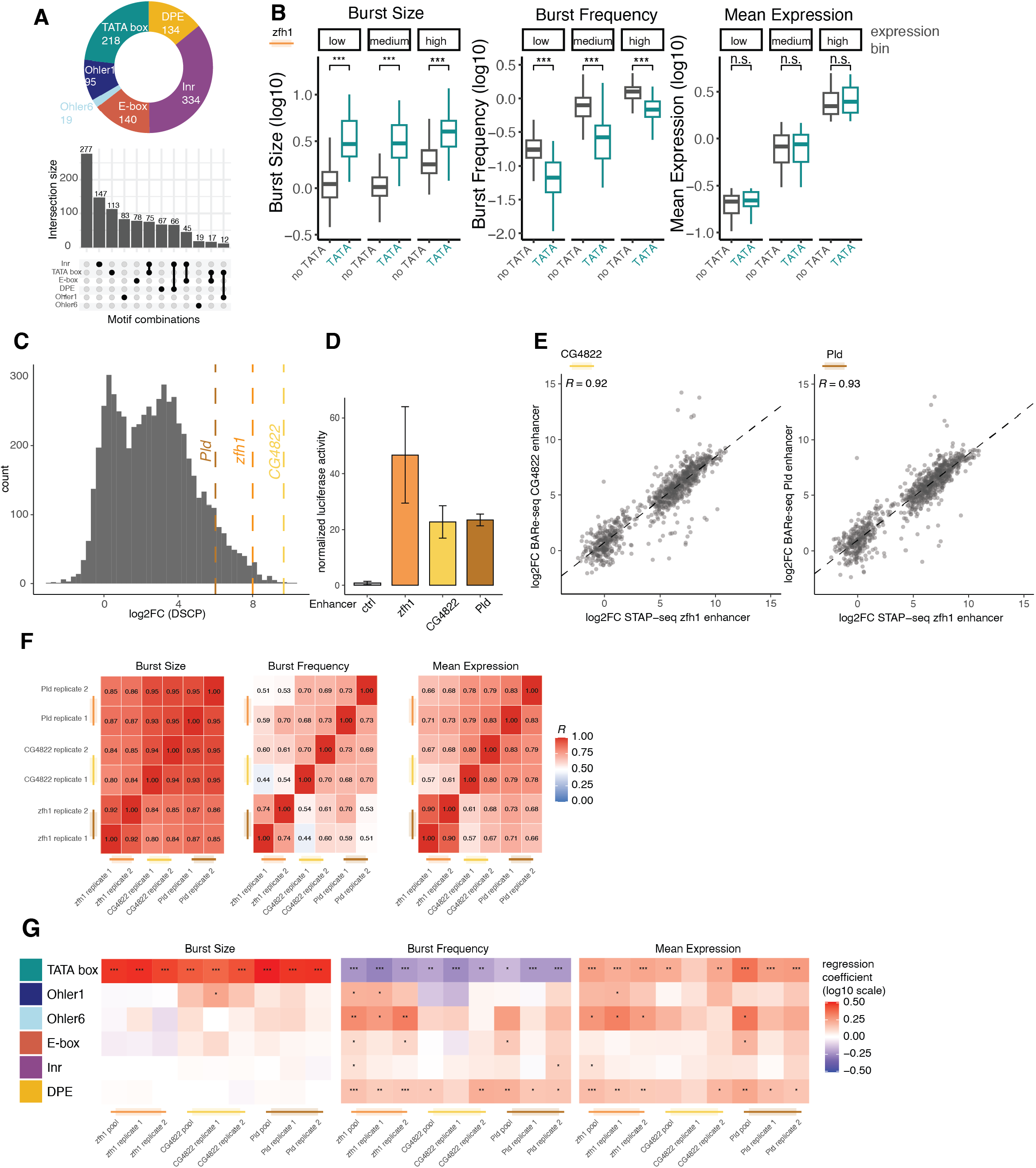
Additional information on the promoter BARe-seq libraries. (A) Promoter motif distribution and upset plot of motif co-occurrence of the selected set of main core promoter motifs in the promoter sequences. (B) Boxplot of grouped comparison of burst parameters of TATA box-containing and non-TATA box promoters in three mean expression bins of similar number of sequences. Wilcoxon rank test Significance markings: (* *p* < 0.05, ** *p* < 0.01, *** *p* < 0.001). (C) Histogram of log_2_ FC values from a published STARR-seq enhancer activity screen (de Almeida et al., 2022). The three enhancers utilized in the promoter-BARe-seq screens are marked by dashed lines. (D) Bar plot showing normalized luciferase activity for the three selected enhancers (zfh1, CG4822, Pld). Error bars represent the standard error of triplicate measurements. (E) Scatter plot of promoter activity derived from BARe-seq in the CG4822 and Pld enhancer contexts compared to published promoter activity data (Haberle et al., 2019). (F) Heatmap showing Pearson’s R values for replicate correlation of individual promoter-BARe-seq experiments prior to pooling. (G) Heatmap showing motif-associated regression coefficients from multivariable regression models of promoter-BARe-seq data in the zfh1, CG4822 and Pld enhancer context. Coefficients are shown on the log_10_-transformed outcome scale before pooling. Significance markings: (* *p* < 0.05, ** *p* < 0.01, *** *p* < 0.001).

**Supplemental figure 4:**
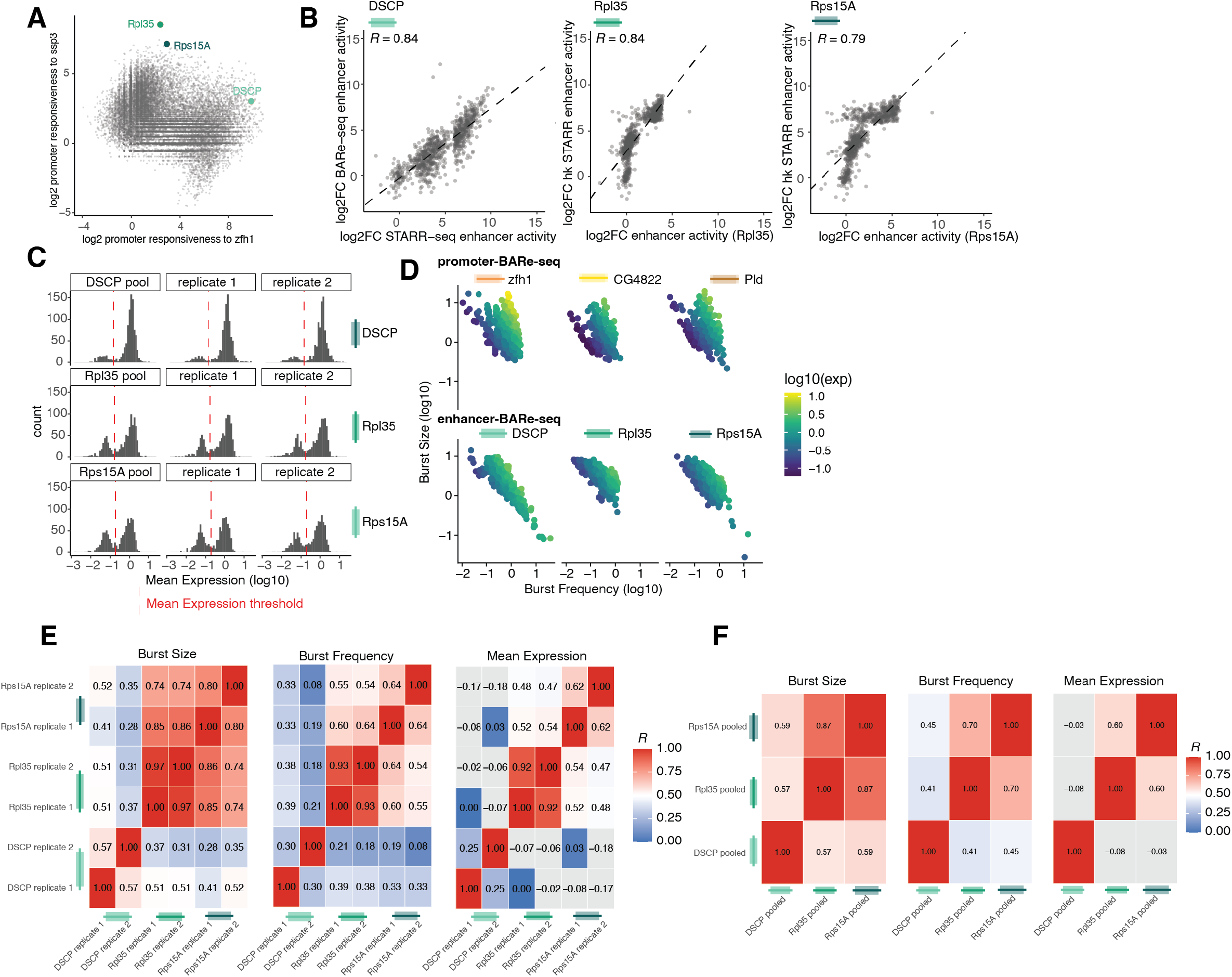
Additional information on the enhancer BARe-seq libraries. (A) Scatter plot of published promoter responsiveness to the developmental zfh1 or the housekeeping ssp3 enhancer (Haberle et al., 2019). The three promoters used with the enhancer-BARe-seq library (DSCP, Rpl35, Rps15A) are highlighted. (B) Scatter plots of enhancer-BARe-seq activities and published enhancer activity values (de Almeida et al., 2022). R = Pearson’s R. (C) Histograms of inferred mean expression. Red dashed lines mark the minimum expression thresholds applied for enhancer BARe-seq filtering (Supplemental Table 3). (D) Scatter plot of burst size as a function of burst frequency, colored by mean expression value. (E) Heatmap showing Pearson’s R values of comparison of replicates of individual enhancer-BARe-seq replicates prior to pooling. (F) Heatmap showing Pearson’s R values of comparison of pooled enhancer-BARe-seq libraries.

**Supplemental Figure 5:**
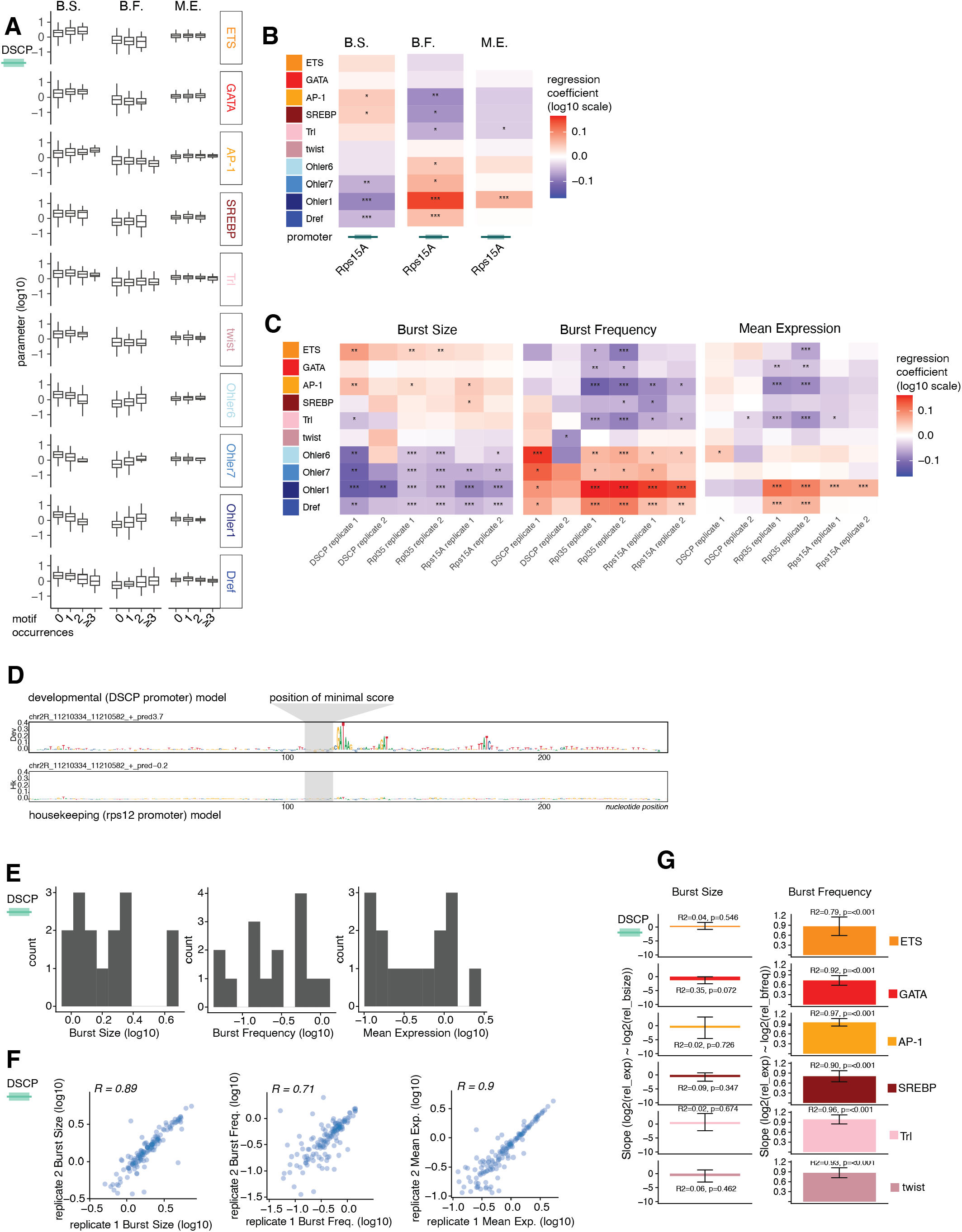
Additional information of the enhancer BARe-seq libraries. (A) Boxplots showing burst parameters aggregated by the number of motif occurrences (x-axis) for the indicated enhancer motifs. B.S. = Burst Size, B.F. = Burst Frequency, M.E. = Mean Expression. (B) Heatmap showing motif-associated regression coefficients from multivariable regression models of enhancer-BARe-seq data in the Rps15A promoter context (log_10_-transformed outcome scale). (C) Heatmap showing motif-associated regression coefficients from multivariable regression models of enhancer-BARe-seq data of individual replicates prior to pooling (log_10_-transformed outcome scale. Significance markings: (* *p* < 0.05, ** *p* < 0.01, *** *p* < 0.001). (D) Schematic showing the strategy for identifying regions of minimal importance (=minimal nucleotide contribution scores (for details see methods)) for motif insertion. (E) Histograms showing the distribution of burst size, burst frequency and mean expression values of unmodified wild-type base enhancers. (F) Scatter plots showing replicate correlations of raw estimated parameters for the motif insertion experiment. (G) Bar graphs of slopes derived from linear regression fits of log_2_-transformed relative expression versus burst parameters. Error bars represent the 95% confidence intervals of the slope estimates. Goodness-of-fit (R^2^) and p-values are indicated for each fit.

## Methods

### Generating libraries

300-bp oligonucleotides containing 133-bp (promoter) or 249-bp (enhancer) candidate sequences were custom synthesized by Twist Biocence. Oligos were amplified and prepared for cloning by sequential PCR steps using KAPA HiFi HotStart ReadyMix (Roche Diagnostics), with bead-based clean ups (AMPure XP Beads, Beckman Coulter A63881) between steps to remove residual primers (Supplemental Figure 1A). All primer sequences are provided in Supplemental Table 1. First starting each oligo pool was amplified with pool-specific primers from 1ng of template DNA within the linear amplification range. Second, plasmid UMIS (pUMIs) were introduced using single-primer linear PCR (50 cycles) using 2.5ng of template per 50μl PCR reaction, with 20 reactions to maintain complexity. Third, Gibson assembly overhangs were added by PCR. PCR cycle numbers were determined empirically to ensure that this amplification was also in a linear range.

The promoter and enhancer of the published reporter vectors for STAP- and STARR-seq assays (Addgene No. 71499 & 86831) were replaced with the promoter and enhancer sequences as summarized in Supplemental table 4 and 6. All reporter backbones were then digested with KpnI and AgeI and gel purified before amplified inserts were cloned by Gibson assembly (New England Biolabs, catalog no. E2611L) using 14 reactions each containing 125ng vector backbone. Assembled libraries were electroporated into MegaX DH10B electrocompetent bacteria (Thermo Fisher ScientificC640003) and expanded in 6L LB-Amp (Luria-Bertani medium plus ampicillin, 100 µg/ml) per library to OD 2.0 and purified with a Qiagen Plasmid *Plus* Mega Kit (catalog no. 12991). Transformation efficiency was monitored by colony counts and plasmid input sequencing. Library representation was assessed by UMI-tagged amplification and sequencing of the plasmid input, as described previously (Neumayr et al. 2019).

### Oligo pool design

#### Promoters

To design the core promoter library, we selected candidate sequences from a previously characterized genome-wide STAP-seq dataset (Haberle et al 2019). Promoter activity values as reported in Haberle et al., 2019 were used to ensure the new library contained robustly active CPs: We selected developmental-specific promoters (log2FC_zfh1> 6 & log2FC_zfh1 - log2FC_ssp3 > 1), housekeeping-specific (log2FC_ssp3 > 5 & log2FC_zfh1 - log2FC_ssp3 < -1), and highly active promoters responsive to both enhancers (log2FC_zfh1> 6 and log2FC_ssp3 > 5). To characterize the sequence architecture of these active promoters, candidates were annotated for the presence of canonical core promoter elements (e.g., TATA box, INR, DPE, Ohler1/6/7, DRE, TCT, and E-box) using the position weight matrix thresholds reported in Haberle et al., 2019. We restricted the pool to the most common specific single- and multi-motif combinations and randomly down-sampled to a maximum of 75 promoters per motif architecture category. We also incorporated 78 random genomic regions with minimal enhancer responsiveness (log2FC_zfh1 & log2FC_ssp3 < 1.1) to serve as negative controls. The selected 133bp promoters were flanked with constant regions including Illumina adapters and a spacer sequence, to generate 300bp oligonucleotides.

#### Enhancers

To assemble the enhancer library, we selected 249-bp enhancer sequences from a previously published synthetic STARR-seq screen (de Almeida et al., 2022). Enhancers were chosen to represent distinct activity profiles based on their enrichment over plasmid library input (log2FC). Developmental enhancers were selected for high developmental activity (log2FC_DSCP > 6) and strong specificity (log2FC_DSCP – log2FC_rps12 > 2). Housekeeping enhancers were filtered for robust housekeeping activity (log2FC_rps12 > 6) and specificity (log2FC_DSCP - log2FC_rps12 < -3). Broadly active enhancers were defined as those highly active in either condition (log2FC > 6) with minimal condition bias (absolute log2FC difference ≤ 1). Sequences were randomly sampled from each class (414 developmental, 400 housekeeping, 14 both, 84 inducible enhancers). Negative controls consisted of 95 inactive genomic regions with low activity in both conditions (log2FC < 1.5). Selected sequences were annotated for motif content using motifmatchr (version 1.18.0, parameters: genome = “dm3”, bg = “genome”, p.cutoff = 5e-04.) and curated PWMs for regulatory motifs in *Drosophila* S2 cells (de Almeida et al., 2022). The 249-bp enhancer sequences were flanked with constant sequences for PCR amplification and cloning to generate 300-bp oligonucleotides.

#### Motif addition

To test the impact of motif addition on enhancer, we designed motif-insertion variants for 14 active enhancers. Candidate insertion sites were selected using base-resolution scores derived from DeepSTARR models trained on developmental and housekeeping enhancer activity (de Almeida 2022). For each 249-bp enhancer, nucleotide contribution scores from both models were summed in a 12-bp sliding window across the sequence to locate the window with the minimum contribution score sum. This region was defined as the most structurally neutral locus for motif insertion. The transcription factor motifs were computationally inserted into the identified 12-bp neutral window. To strictly preserve the 249-bp enhancer length, shorter motifs were centered and padded with the original endogenous sequence of the window, while longer motifs were symmetrically centered around the window’s midpoint. Motif sequences and insertion coordinates are listed in Supplemental Table 8.

All genomic coordinates refer to the dm3 assembly of the *Drosophila* melanogaster genome. Sequence manipulation was performed in R using seqinr (v4.2-8) and stringr (v1.5.0). Genomic coordinates and oligonucleotide DNA sequences for all libraries are provided in Supplemental Tables 4 and 6.

### Cell culture

Drosophila S2 cells (Thermofisher R69007) were grown in Schneider’s Insect medium (Gibco 21720001) with 10% fetal bovine serum (FBS) (Sigma-Aldrich, heat inactivated) and 1% P/S (Gibco 15140122) at 27°C and 0.4% CO_2_.

### Dual luciferase assay

To validate enhancers used for STAP-seq, a dual firefly and Renilla luciferase reporter assay was used, reading out luciferase activity as described previously (Arnold et al., 2013). Firefly luciferase reporter plasmids were based on a pGL3 luciferase reporter vector (Promega, E1751), in which the SV40 promotor was replaced by the synthetic *Drosophila* DSCP. The DNA sequences to be tested as enhancers were PCR-amplified from genomic DNA and cloned upstream of the DSCP (using KpnI). Each luciferase experiment was performed in three biological replicates, transfecting 60’000 S2 cells with 100ng of the firefly luciferase reporter and 5ng of a *Renilla* luciferase expression vector (pRL-ubi63E Addgene #74280) (Arnold et al., 2013) with FuGENE HD Transfection Reagent (Promega, E2311). Firefly and Renilla luciferase activity were measured 48h after transfection in a BioTek Synergy H1 Plate Reader (Agilent) using the Dual Luciferase Reporter Assay Kit (Promega, E1960) and normalized to *Renilla* luciferase activity to control for cell number variability and transfection efficiency.

### Electroporations

Cells were seeded at 3*10^6/ml density the day before electroporation. Per library we performed at least 2 replicate experiments. For each of the biological replicate, 50*10^6 cells cells were resuspended in 100 µL of a 1:1 dilution of HyClone MaxCyte electroporation buffer and serum-free Schneider’s and electroporated with 5 µg of the input libraries (see previous section) using the MaxCyte-STX system (‘Optimization 1’ protocol). Cells were then incubated in DNase I (Worthington Biochemical, 2000 U/mL) for 30 min and harvested for RNA extraction 24 h after addition of culture media.

### Screens

Cells were counted and 1-2 million cells were processed per replicate. RNA was extracted (Quiagen RNA mini kit, Catalogue no. 74104) and processed following a downscaled and optimized version of the previously described UMI-STARR-seq protocol or STAP-seq protocol (Neumayer et al. 2019, Haberle et al. 2019). In brief both protocols contain a reporter specific reverse transcription step and an additional step for the introduction of a tUMI - either by RNA adapter annealing for promoter reporter transcripts or by linear single primer PCR following reverse transcription for enhancer reporter transcripts. The resulting transcripts preserve the association between each reporter sequence, its pUMI and its tUM, allowing transcript counts to be assigned to individual reporter alleles.

### Illumina sequencing

Next-generation sequencing was performed at the VBCF NGS facility on a HiSeq 2000 or NovaSeq SP platform (36bp paired-end or 50/150bp paired-end), following the manufacturer’s protocol. For promoter libraries the ligated 5’ RNA adapters contained a 4nt sample barcode and the 10nt tUMI, while for enhancer libraries we used standard Illumina i5 indexes as well as tUMIs at the i7 index. In both cases the pUMI was part of the reverse read together with the promoter/enhancer sequence.

### MPRA data analysis

A custom index of the designed oligonucleotide sequences of each library was generated using the buildindex function from the Rsubread R package version 2.10.0 (Liao et al 2019). Paired-end reads were aligned using the align function using type = “dna”, unique = TRUE, maxMismatches = 1 for 36 or 50bp reads and 3 for 150bp sequencing reads, nTrim = 1. Oligos with coverage below 2% of the median library cpverage and reads containing poly-G stretches longer than 10bp were discarded. To reduce overcounting from sequencing errors, pUMIs and transcript UMIs (tUMIs) were collapsed sequentially. Reads were first grouped by oligo identifier, and exact pUMI– tUMI combinations were deduplicated. Within each oligo, pUMIs were sorted by RNA count, and lower-abundance pUMIs within a Levenshtein distance of three from a higher-abundance pUMI were collapsed using stringdist version 0.9.8. After pUMI correction, tUMIs were collapsed using a Hamming distance of one, as described previously (de Almeida et al., 2022). After collapsing we discarded reads where the fixed part of the pUMI did not match the expected sequence structure and then generated tables of transcript counts per plasmid, which were subsequently used for burst parameter inference. Activity scores for each oligo were estimated as log2 fold-change of RNA counts relative to plasmid DNA input using DESeq2 (Love et al., 2014), following de Almeida et al. (2022).

### Modeling of burst parameters: Two-state model of transcriptional bursting

We modeled allele-resolved reporter transcript counts using a two-state statistical model of transcriptional bursting proposed by Peccoud and Ycart (Peccoud & Ycart, 1995). In this model, each allele switches between ON and OFF states with activation and inactivation rates *k*_*on*_ and *k*_*off*_. Transcripts are synthesized in the ON state at a constant rate of synthesis *k*_*syn*_ and degraded with a constant rate *k*_*deg*_. At steady state, the number of observed RNA molecules *X* are captured by the chemical master equation with the constants of activation, deactivation and synthesis normalized by the rate of degradation (*k*_*on*_/*k*_*deg*_, *k*_*off*_/*k*_*deg*_, *k*_*syn*_/*k*_*deg*_) (Peccoud & Ycart, 1995). For convention, the *k*_*deg*_ term is removed from the notation, yet the kinetic constant values are considered as the corresponding ratios with respect to *k*_*deg*_.

In the bursty limit where *k*_*off*_ ≫ *k*_*on*_ and *k*_*off*_ > *k*_*deg*_, the steady-state distribution of transcript counts can be approximated by a Gamma-Poisson distribution, equivalent to a negative binomial distribution (*NB*) (Grün et al., 2014; Raj et al., 2006; Shahrezaei & Swain, 2008)

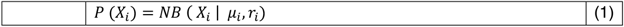

where *X*_*i*_ is the number of transcripts observed for allele *i*. *μ*_*i*_ is the mean and *r*_*i*_ is the dispersion parameter of the *NB*. In terms of kinetic parameters, *μ* and *r* are equivalent to (Grün et al., 2014)

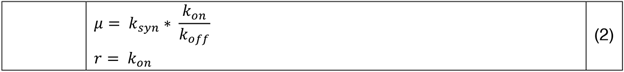

Burst parameters (frequency and size) can be inferred from the kinetic parameters (Kim & Marioni, 2013; Raj et al., 2006)

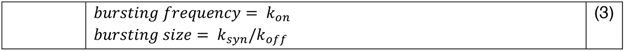

The burst parameters can also be expressed in terms of the negative binomial parameters as follows(Raj et al., 2006; Grün et al., 2014)

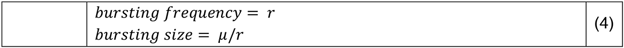

### Implementation

Because alleles with zero captured transcripts are not observed in BARe-seq, transcript count distributions were modeled with a *zero* − *truncated NB*:

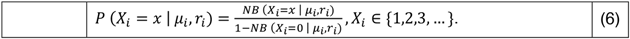

For each regulatory sequence, the set of parameters *θ*_*i*_ = (*μ*_*i*_, *r*_*i*_) were inferred by maximum likelihood estimation:

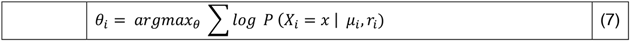

Optimization was performed in R using optim with the BFGS method, starting values (1,1) and a maximum of 10000 iterations.

### Kolmogorov-Smirnov test

Goodness of fit was assessed by comparing the relative frequency of observed data counts versus the theoretical probability distribution estimated with the inferred parameters under the fitted 0-truncated NB model using the Kolmogorov-Smirnov test with the help of the ks.test in R. P values are corrected for multiple testing using the Benjamini–Hochberg method and sequences with P adj > 0.01 were retained.

### Mean expression threshold selection

The inferred parameters showed a subset of data with very low *μ* and *r* parameter values. These estimates were interpreted as corresponding to regulatory elements with low transcriptional activity, based on the low number of captured transcript counts. Such observations were excluded from subsequent analyses, since they were more unstable in inference. Thresholds for *μ* were defined as the local minimum between the two modes of its distribution (i.e. the point of lowest density). Dataset-specific thresholds are listed in Supplemental Table 3.

### Data simulation

We performed a simulation of 5000 sequences to explore which regions of the parameter space are confidently estimated. We sampled random transcript counts from a *zero* − *truncated NB* distribution with different combinations of *μ* and *r* parameter values generously covering the range of inferred values of the experimental data. *μ* ranged from 10^-2.85^ to 10^1.15^ and *r* from 10^-2.85^ to 10^2.25^. To guarantee that the parameter inference is not affected by a low number of observations, we drew a high number of allele-level transcript counts (i.e. 1700) per simulated sequence.

### Multivariable analysis

Associations between motif composition and the three burst parameters were quantified using multivariable linear regression. For each dataset and burst parameter - log10-transformed burst size, burst frequency or mean expression was modeled as a function of motif counts. While promoter motifs were either absent or only present once per sequence, motif counts in enhancers were clipped at two or three occurrences (Supplemental Figure 5) before fitting to reduce the influence of high-count outliers. All selected motifs were included simultaneously in the regression model using lm in R. Regression coefficients therefore represent conditional associations between each motif and the readout, controlling for the other motifs included in the model.

### Meta-analysis

For each motif–outcome combination, regression coefficients and standard errors from the multivariable linear models were pooled across datasets (promoters in three enhancer contexts) using inverse-variance weighted fixed-effect meta-analysis using the R metafor package version 3.4-0. Only motif–outcome combinations represented in all three datasets were included. Pooled estimates were reported on the log10 scale and converted to fold changes for interpretation; 95% confidence intervals and p-values were derived from the pooled meta-analysis. Motifs were classified as significant if the pooled association reached p<0.05 for either burst size or burst frequency.

### Statistics and data visualization

All statistical calculations and graphical displays were performed in R statistical computing environment (v.4.2.0) and using the R packages data.table (v1.14.2), dplyr (version 1.1.2) and ggplot2 (v.3.4.2). Unless stated otherwise, in all box plots, the central line denotes the median, the box encompasses 25th to 75th percentile (interquartile range) and the whiskers extend to 1.5× interquartile range.

## Data availability

The raw sequencing data are available from GEO (https://www.ncbi.nlm.nih.gov/geo/) under accession number GSE342901.

## Code availability

Custom scripts to analyze the data and plot the figures are available on GitHub: https://github.com/florbeer/BARe-seq.git

## Tables

1. Library cloning information
2. Information on experimental batches, replicates and pooling
3. Mean expression thresholds
4. Meta data of promoter sequences
5. Inferred burst parameters of promoter BARe-seq
6. Meta data of enhancer sequences
7. Inferred burst parameters of enhancer BARe-seq
8. Meta data of enhancers with pasted motifs
9. Inferred burst parameters of motif insertions in enhancers

## References

Arnold, C. D., Gerlach, D., Stelzer, C., Boryń, Ł. M., Rath, M., & Stark, A. (2013). Genome-Wide Quantitative Enhancer Activity Maps Identified by STARR-seq. Science, 339(6123), 1074–1077. 10.1126/science.1232542

Arnold, C. D., Zabidi, M. A., Pagani, M., Rath, M., Schernhuber, K., Kazmar, T., & Stark, A. (2017). Genome-wide assessment of sequence-intrinsic enhancer responsiveness at single-base-pair resolution. Nature Biotechnology, 35(2), 136–144. 10.1038/nbt.3739

Bergman, D. T., Jones, T. R., Liu, V., Ray, J., Jagoda, E., Siraj, L., Kang, H. Y., Nasser, J., Kane, M., Rios, A., Nguyen, T. H., Grossman, S. R., Fulco, C. P., Lander, E. S., & Engreitz, J. M. (2022). Compatibility rules of human enhancer and promoter sequences. Nature, 607(7917), 176–184. 10.1038/s41586-022-04877-w

Bothma, J. P., Garcia, H. G., Esposito, E., Schlissel, G., Gregor, T., & Levine, M. (2014). Dynamic regulation of eve stripe 2 expression reveals transcriptional bursts in living Drosophila embryos. Proceedings of the National Academy of Sciences of the United States of America, 111(29), 10598–10603. 10.1073/pnas.1410022111

de Almeida, B. P., Reiter, F., Pagani, M., & Stark, A. (2022). DeepSTARR predicts enhancer activity from DNA sequence and enables the de novo design of synthetic enhancers. Nature Genetics, 54(5), Article 5. 10.1038/s41588-022-01048-5

Elowitz, M. B., Levine, A. J., Siggia, E. D., & Swain, P. S. (2002). Stochastic Gene Expression in a Single Cell. Science, 297(5584), 1183–1186. 10.1126/science.1070919

Falo-Sanjuan, J., Lammers, N. C., Garcia, H. G., & Bray, S. J. (2019). Enhancer Priming Enables Fast and Sustained Transcriptional Responses to Notch Signaling. Developmental Cell, 50(4), 411–425.e8. 10.1016/j.devcel.2019.07.002

Fukaya, T., Lim, B., & Levine, M. (2016). Enhancer Control of Transcriptional Bursting. Cell, 166(2), 358–368. 10.1016/j.cell.2016.05.025

Garcia, H. G., Tikhonov, M., Lin, A., & Gregor, T. (2013). Quantitative Imaging of Transcription in Living *Drosophila* Embryos Links Polymerase Activity to Patterning. Current Biology, 23(21), 2140–2145. 10.1016/j.cub.2013.08.054

Golding, I., Paulsson, J., Zawilski, S. M., & Cox, E. C. (2005). Real-Time Kinetics of Gene Activity in Individual Bacteria. Cell, 123(6), 1025–1036. 10.1016/j.cell.2005.09.031

Grün, D., Kester, L., & van Oudenaarden, A. (2014). Validation of noise models for single-cell transcriptomics. Nature Methods, 11(6), Article 6. 10.1038/nmeth.2930

Haberle, V., Arnold, C. D., Pagani, M., Rath, M., Schernhuber, K., & Stark, A. (2019). Transcriptional cofactors display specificity for distinct types of core promoters. Nature, 570(7759), Article 7759. 10.1038/s41586-019-1210-7

Hendy, O., Campbell, J., Weissman, J. D., Larson, D. R., Singer, D. S., & Tansey, W. P. (2017). Differential context-specific impact of individual core promoter elements on transcriptional dynamics. Molecular Biology of the Cell, 28(23), 3360–3370. 10.1091/mbc.e17-06-0408

Hoppe, C., Bowles, J. R., Minchington, T. G., Sutcliffe, C., Upadhyai, P., Rattray, M., & Ashe, H. L. (2020). Modulation of the Promoter Activation Rate Dictates the Transcriptional Response to Graded BMP Signaling Levels in the Drosophila Embryo. Developmental Cell, 54(6), 727–741.e7. 10.1016/j.devcel.2020.07.007

Hornung, G., Bar-Ziv, R., Rosin, D., Tokuriki, N., Tawfik, D. S., Oren, M., & Barkai, N. (2012). Noise–mean relationship in mutated promoters. Genome Research, 22(12), 2409–2417. 10.1101/gr.139378.112

Keller, S. H., Jena, S. G., Yamazaki, Y., & Lim, B. (2020). Regulation of spatiotemporal limits of developmental gene expression via enhancer grammar. Proceedings of the National Academy of Sciences of the United States of America, 117(26), 15096–15103. 10.1073/pnas.1917040117

Kim, J. K., & Marioni, J. C. (2013). Inferring the kinetics of stochastic gene expression from single-cell RNA-sequencing data. Genome Biology, 14(1), R7. 10.1186/gb-2013-14-1-r7

Kwasnieski, J. C., Fiore, C., Chaudhari, H. G., & Cohen, B. A. (2014). High-throughput functional testing of ENCODE segmentation predictions. Genome Research, 24(10), 1595–1602. 10.1101/gr.173518.114

Larsson, A. J. M., Johnsson, P., Hagemann-Jensen, M., Hartmanis, L., Faridani, O. R., Reinius, B., Segerstolpe, Å., Rivera, C. M., Ren, B., & Sandberg, R. (2019). Genomic encoding of transcriptional burst kinetics. Nature, 565(7738), Article 7738. 10.1038/s41586-018-0836-1

Larsson, A. J. M., Ziegenhain, C., Hagemann-Jensen, M., Reinius, B., Jacob, T., Dalessandri, T., Hendriks, G.-J., Kasper, M., & Sandberg, R. (2021). Transcriptional bursts explain autosomal random monoallelic expression and affect allelic imbalance. PLoS Computational Biology, 17(3), e1008772. 10.1371/journal.pcbi.1008772

Lee, C., Shin, H., & Kimble, J. (2019). Dynamics of Notch-Dependent Transcriptional Bursting in Its Native Context. Developmental Cell, 50(4), 426–435.e4. 10.1016/j.devcel.2019.07.001

Lim, B., Fukaya, T., Heist, T., & Levine, M. (2018). Temporal dynamics of pair-rule stripes in living Drosophila embryos. Proceedings of the National Academy of Sciences, 115(33), 8376–8381. 10.1073/pnas.1810430115

Nicolas, D., Phillips, N. E., & Naef, F. (2017). What shapes eukaryotic transcriptional bursting? Molecular BioSystems, 13(7), 1280–1290. 10.1039/C7MB00154A

Ochiai, H., Hayashi, T., Umeda, M., Yoshimura, M., Harada, A., Shimizu, Y., Nakano, K., Saitoh, N., Liu, Z., Yamamoto, T., Okamura, T., Ohkawa, Y., Kimura, H., & Nikaido, I. (2020). Genome-wide kinetic properties of transcriptional bursting in mouse embryonic stem cells. Science Advances, 6(25), eaaz6699. 10.1126/sciadv.aaz6699

Ohler, U. (2006). Identification of core promoter modules in Drosophila and their application in accurate transcription start site prediction. Nucleic Acids Research, 34(20), 5943–5950. 10.1093/nar/gkl608

Peccoud, J., & Ycart, B. (1995). Markovian Modeling of Gene-Product Synthesis. Theoretical Population Biology, 48(2), 222–234. 10.1006/tpbi.1995.1027

Pennington, K. L., Marr, S. K., Chirn, G.-W., & Marr, M. T. (2013). Holo-TFIID controls the magnitude of a transcription burst and fine-tuning of transcription. Proceedings of the National Academy of Sciences, 110(19), 7678–7683. 10.1073/pnas.1221712110

Pfeiffer, B. D., Jenett, A., Hammonds, A. S., Ngo, T.-T. B., Misra, S., Murphy, C., Scully, A., Carlson, J. W., Wan, K. H., Laverty, T. R., Mungall, C., Svirskas, R., Kadonaga, J. T., Doe, C. Q., Eisen, M. B., Celniker, S. E., & Rubin, G. M. (2008). Tools for neuroanatomy and neurogenetics in Drosophila. Proceedings of the National Academy of Sciences, 105(28), 9715–9720. 10.1073/pnas.0803697105

Pimmett, V. L., Dejean, M., Fernandez, C., Trullo, A., Bertrand, E., Radulescu, O., & Lagha, M. (2021). Quantitative imaging of transcription in living Drosophila embryos reveals the impact of core promoter motifs on promoter state dynamics. Nature Communications, 12(1), Article 1. 10.1038/s41467-021-24461-6

Raj, A., Peskin, C. S., Tranchina, D., Vargas, D. Y., & Tyagi, S. (2006). Stochastic mRNA Synthesis in Mammalian Cells. PLOS Biology, 4(10), e309. 10.1371/journal.pbio.0040309

Raj, A., & van Oudenaarden, A. (2008). Nature, nurture, or chance: Stochastic gene expression and its consequences. Cell, 135(2), 216–226. 10.1016/j.cell.2008.09.050

Rosales-Alvarez, R. E., Rettkowski, J., Herman, J. S., Dumbović, G., Cabezas-Wallscheid, N., & Grün, D. (2023). VarID2 quantifies gene expression noise dynamics and unveils functional heterogeneity of ageing hematopoietic stem cells. Genome Biology, 24(1), 148. 10.1186/s13059-023-02974-1

Sanchez, A., & Golding, I. (2013). Genetic Determinants and Cellular Constraints in Noisy Gene Expression. Science, 342(6163), 1188–1193. 10.1126/science.1242975

Senecal, A., Munsky, B., Proux, F., Ly, N., Braye, F. E., Zimmer, C., Mueller, F., & Darzacq, X. (2014). Transcription factors modulate c-Fos transcriptional bursts. Cell Reports, 8(1), 75–83. 10.1016/j.celrep.2014.05.053

Serebreni, L., Pleyer, L.-M., Haberle, V., Hendy, O., Vlasova, A., Loubiere, V., Nemčko, F., Bergauer, K., Roitinger, E., Mechtler, K., & Stark, A. (2023). Functionally distinct promoter classes initiate transcription via different mechanisms reflected in focused versus dispersed initiation patterns. The EMBO Journal, 42(10), e113519. 10.15252/embj.2023113519

Shahrezaei, V., & Swain, P. S. (2008). Analytical distributions for stochastic gene expression. Proceedings of the National Academy of Sciences, 105(45), 17256–17261. 10.1073/pnas.0803850105

Tunnacliffe, E., & Chubb, J. R. (2020). What Is a Transcriptional Burst? Trends in Genetics, 36(4), 288–297. 10.1016/j.tig.2020.01.003

van Arensbergen, J., FitzPatrick, V. D., de Haas, M., Pagie, L., Sluimer, J., Bussemaker, H. J., & van Steensel, B. (2017). Genome-wide mapping of autonomous promoter activity in human cells. Nature Biotechnology, 35(2), 145–153. 10.1038/nbt.3754

Yáñez-Cuna, J. O., Arnold, C. D., Stampfel, G., Boryń, Ł. M., Gerlach, D., Rath, M., & Stark, A. (2014). Dissection of thousands of cell type-specific enhancers identifies dinucleotide repeat motifs as general enhancer features. Genome Research, 24(7), 1147–1156. 10.1101/gr.169243.113

Yokoshi, M., Kawasaki, K., Cambón, M., & Fukaya, T. (2022). Dynamic modulation of enhancer responsiveness by core promoter elements in living Drosophila embryos. Nucleic Acids Research, 50(1), 92–107. 10.1093/nar/gkab1177

Zabidi, M. A., Arnold, C. D., Schernhuber, K., Pagani, M., Rath, M., Frank, O., & Stark, A. (2015). Enhancer–core-promoter specificity separates developmental and housekeeping gene regulation. Nature, 518(7540), 556–559. 10.1038/nature13994

